# Host regulator PARP1 contributes to sex differences and immune responses in a mouse model of tuberculosis

**DOI:** 10.1101/2021.04.21.440820

**Authors:** Stefanie Krug, Alvaro A. Ordonez, Mariah Klunk, Bong Gu Kang, Sanjay K. Jain, Ted M. Dawson, Valina L. Dawson, William R. Bishai

## Abstract

Tuberculosis (TB) is a devastating infectious disease responsible for nearly 2 million deaths annually that has a poorly understood male bias. Elucidating the basis of this male bias may enable precision medicine interventions for TB treatment and prevention. Here, we identify the master regulator Poly(ADP-ribose) Polymerase 1 (PARP1) as a driver of TB sex differences. We found that infection with *M. tuberculosis* (*M. tb*) triggers robust PARP activation in mouse lungs, suggesting that PARP1 activation is a fundamental host response to TB. Remarkably, PARP1 deletion abolished known sex differences in TB cytokine responses and blunted the early induction of TNFα, IL-1ß, IFNγ, MCP-1, and IL-6, particularly in male mice. In contrast, PARP1 was required for IL-10 induction in male or female mice. PARP1 deletion was protective against TB in female mice, resulting in significantly prolonged survival and reduced bacterial burden, but impaired TB containment in male mice. Our findings indicate that PARP1 contributes to TB sex differences via sexually divergent immune regulation and uniquely enhances early proinflammatory responses in males that prove beneficial for TB containment.

## Introduction

Tuberculosis (TB) is a devastating global health threat responsible for over 1.5 million deaths every year (1). TB has an appreciated male bias, with nearly twice as many adult men developing active TB as women (2–4). Moreover, TB mortality, lesion severity, cavitation rates and treatment responses tend to be worse in men than in women (5–8). A male bias in disease severity and lethality has also been demonstrated in mice infected with *Mycobacterium tuberculosis* (*M. tb*) and non-tuberculous mycobacteria, including *M. marinum* and *M. avium* (9–12). While these patterns appear to be driven primarily by biological factors, surprisingly little is known about genetic and hormonal factors which underlie this male bias (4, 13).

The eukaryotic master regulator Poly(ADP-ribose) Polymerase-1 (PARP1) has emerged as a critical cofactor of NF-κB activation and TNFα signaling but its role in TB has not yet been studied (14–18). PARP1 is a nuclear enzyme that regulates fundamental cellular functions by modifying nuclear proteins, including itself, and sites of DNA damage post-translationally with poly(ADP-ribose) polymers (PAR) in a process termed PARylation (15, 19–22). PARylation is a potent, multi-faceted stress response element that can trigger increased proinflammatory gene expression, DNA repair, metabolic changes, and cell death (15, 23, 24). PARP1 also controls numerous immune functions, including immune cell activation, differentiation and recruitment; NF-κB signaling; and the production of TNFα, IL-6, IL-1ß, IFNγ, MCP-1 and nitric oxide (NO) (14, 23, 25). Furthermore, PARP1 amplifies and sustains inflammation by inducing proinflammatory mediators and oxidative stress conditions, which in turn continue to stimulate PARP1 activity in a positive-feedback loop (23, 25). Consequently, PARP1 has been connected to inflammatory disorders, including endotoxic shock, sepsis, asthma, COPD and ARDS, and PARP inhibition reduces inflammation and disease severity in numerous inflammatory conditions (23–26). Since PARP1 regulates key processes that are important for TB containment as well as TB immunopathology, we hypothesized that PARP1 might be a core component of the host response to TB.

PARP1 is also a regulator of biological sex differences, including in stress resistance, inflammation and aging (27–29). PARP1 is involved in silencing the inactive X chromosome, which contains genes involved in pathogen recognition and cytokine signaling, sex hormone signaling, and sexually divergent cell death mechanisms (27, 30–34). Consequently, sexually divergent effects of PARP1 deletion or inhibition have been reported in cell death, brain injury, inflammation, nephrotoxicity, carcinogenesis and aging (29, 30, 32, 34–38). Therefore, PARP1 might have distinct functions in the host response to TB in men and women and similarly contribute to sexually divergent immune regulation in TB infection.

We therefore investigated whether PARP1 may regulate the TB host response, inflammation and sex effects. Here, we demonstrate that PARP1 is activated during *M. tb* infection in mice and serves as a driver of TB inflammation and lung damage. Interestingly, PARP1 deficiency in knockout mice reduced elevated proinflammatory responses in male WT mice to female levels but was only protective in female mice. Our results indicate that PARP1 is an important driver of TB sex differences; in males it uniquely enhances early proinflammatory cytokine production resulting in improved male TB containment.

## Results

### TB infection activates PARP1 in mouse lungs

PARP1 is a host enzyme that regulates cellular functions by post-translationally modifying its targets with poly(ADP-ribose) polymers (PAR) (15). Since these polymers are rapidly degraded, PARP1 activity can be assessed by quantifying PAR levels, which appear as high molecular weight smears on a PAR immunoblot with a pronounced focus around 116 kDa resulting from PARP1 automodification (24, 39). To determine if PARP1 is activated during TB infection, we first evaluated PAR levels in the lungs of age-matched uninfected or *M. tb*-infected C3HeB/FeJ mice (**Fig. 1a-b**). While PAR levels were typically low in uninfected lungs, we detected robust PAR signals in infected lungs that increased over the course of infection, with levels 20- and 80-fold higher than in uninfected lungs (*p*<0.02) at 60 and 120 days post infection (dpi), respectively (**Fig. 1a-b**).

**Figure 1.**
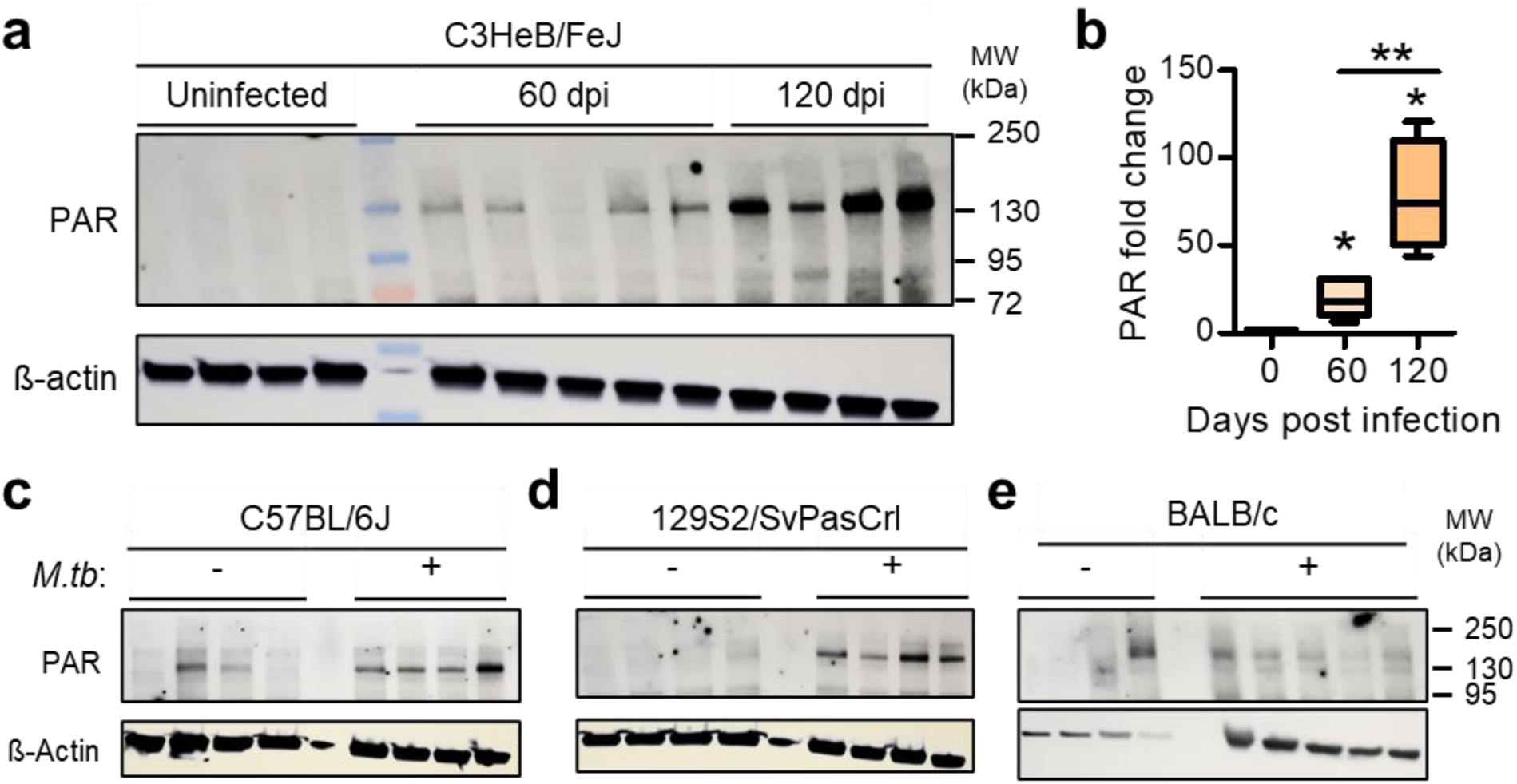
TB infection triggers PARP activation in mouse lungs. (a) Representative PAR immunoblot and (b) densitometric analysis of lung PAR levels in uninfected and *M. tb*-infected C3HeB/FeJ mice 60 and 120 days post infection (dpi) with aerosolized *M. tb* H37Rv (lung implantation: 2.74 log_10_ CFU). Each lane represents an individual mouse. Uninfected controls matched the age of mice at 60 dpi. Box plots illustrate PAR intensity relative to the uninfected average after normalizing to ß-actin (n=4-5). The central line represents the median, the box extends from the 25^th^ to 75^th^ percentiles, and the whiskers represent the range (min to max). Mean PAR intensity in infected lungs is 19.92-fold and 78.34-fold that of uninfected lungs at 60 and 120 dpi, respectively. *\*, p =* 0.016 by unpaired t-test with Welch’s correction for unequal variance. *\*\*, p* = 0.0059 by unpaired t test. (c-d) Representative PAR immunoblots showing lung PAR in age-matched uninfected or *M. tb*-infected (c) C57BL/6J, (d) 129S2/SvPasCrl or (e) BALB/c mice. Each lane represents an individual mouse. C57BL/6J and 129S2/SvPasCrl mice were infected for 51 days (lung implantation: 1.85 log_10_ CFU) and BALB/c mice for 60 days (lung implantation: 2.74 log_10_ CFU). PAR levels are consistently elevated in infected compared to uninfected lungs.

We next assessed the PAR response to infection in three additional mouse strains routinely used to model TB: C57BL/6, 129S2, and BALB/c mice (**Fig. 1c-e**). All strains contained bacterial numbers to around 6 log_10_ CFUs in the lung following low-dose aerosol infection, with no differences in bacterial burden and modest pathology after 7 weeks of infection (not shown). Despite variable baseline PARP1 activity in uninfected lungs, *M. tb* infection consistently induced PAR formation in all three strains, suggesting that PARP1 activation might be a conserved host response to TB. Importantly, treatment with the PARP inhibitor talazoparib (Tp) reduced PAR formation in *M. tb*-infected lungs in a dose-dependent manner, indicating that *M. tb*-induced PARP1 activation can be reversed pharmacologically (**Fig. 2**). Therefore, PARP1 activation appears to be a core component of the host response to *M. tb.* infection and may play a critical host regulatory role in TB.

**Fig. 2:**
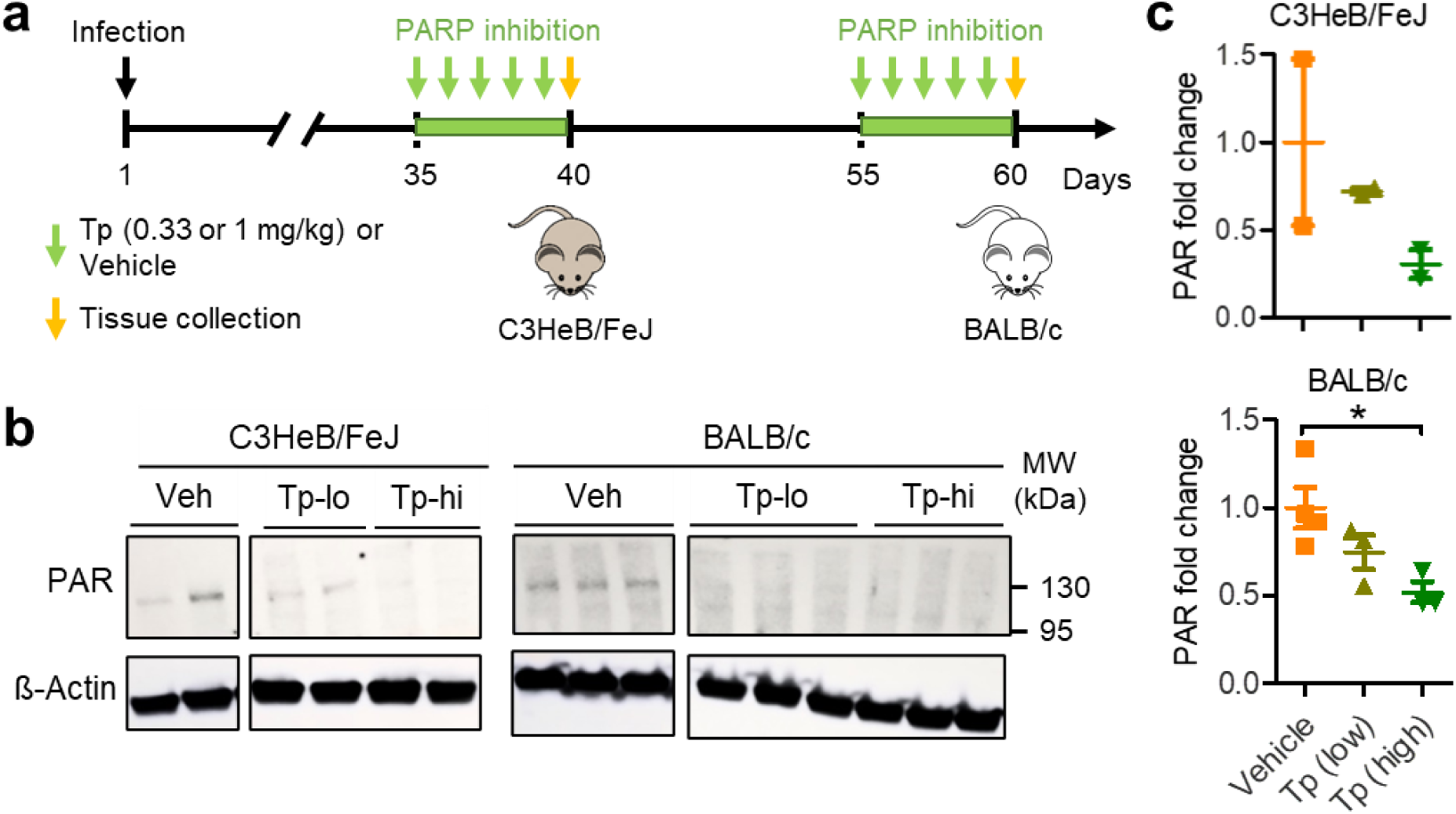
PARP inhibitor talazoparib reduces TB-induced PAR formation in mouse lungs. (a) Schematic overview of study assessing the effects of short-course PARP inhibition on TB-induced PAR formation. Female C3HeB/FeJ or BALB/c mice (10-12 weeks old) were aerosol-infected with *M. tb* H37Rv (implantation: 2.95 log_10_ CFUs) and treated with the PARP1/2 inhibitor talazoparib or vehicle for 5 consecutive days by oral gavage before the lungs were collected for PAR analysis. C3HeB/FeJ mice were treated from 35 to 40 dpi and BALB/c mice from 55 to 60 dpi. (b) Representative PAR immunoblots and (c) densitometric analyses of lung PAR levels in TB-infected C3HeB/FeJ (40 dpi) and BALB/c mice (60 dpi) after 5 days of vehicle (Veh; 264 µl/kg DMSO), low dose (lo; 0.33 mg/kg) or high dose (hi; 1 mg/kg) talazoparib (Tp) administration by oral gavage. Each lane and data point correspond to an individual mouse. PAR levels were normalized to ß-actin and are expressed relative to the average intensity in vehicle-treated mice. *, *p* = 0.0231 by unpaired t-test. The PARP1/2 inhibitor talazoparib reduced lung PAR levels in *M. tb*-infected mice in a dose-dependent manner.

### PARP1 promotes proinflammatory immune responses in male but not in female mice

To characterize the function of PARP1 in TB, we challenged PARP1-deficient 129S-*Parp1^tm1Zqw^*/J (PARP1^-/-^) and control 129S1/SvlmJ (WT) mice with *M. tb* H37Rv via the aerosol route and evaluated immune responses, disease progression, inflammation and survival. In 129S-*Parp1^tm1Zqw^*/J mice, exon 2 of PARP1 is disrupted, and no enzymatic PARP1 activity is detectable in homozygous mutants (40). These PARP1-deficient mice are viable, fertile, and display no physical or behavioral abnormalities (28). We routinely verified the genotype of PARP1^-/-^ mice by PCR and confirmed the absence of PARP1 protein by Western blot (**Supplementary Fig. S1**).

Since PARP1 regulates several immune signaling networks that are critical in the TB host response, we hypothesized that PARP1^-/-^ mice would demonstrate altered immune responses to *M. tb* infection (14, 23, 41). After confirming that uninfected WT and PARP1^-/-^ mice produce comparable lung levels of cytokines and chemokines (**Supplementary Fig. S2**), we next compared the induction of these factors in male and female WT and PARP1^-/-^ mice over the course of infection. Dibbern *et al.* previously reported that male C57BL/6 mice produce higher levels of proinflammatory cytokines one month after *M. tb* infection than female mice (9). Consistent with their findings, we observed markedly higher induction of the proinflammatory cytokines and chemokines, TNFα, IFNγ, IL-6, IL-1β and MCP-1, in 129S1 WT males compared to WT females by 28 dpi (**Fig. 3a**). Remarkably, PARP1 deletion abolished these sex differences and reduced the levels of proinflammatory cytokines in male mice at 28 dpi to female levels (**Fig. 3b**). In contrast, lung levels of IL-2, IL-12, IL-17 and MIP-1α were largely comparable between uninfected and infected mice and appeared to be independent of PARP1 or sex (**Supplementary Fig. S3**).

**Figure 3.**
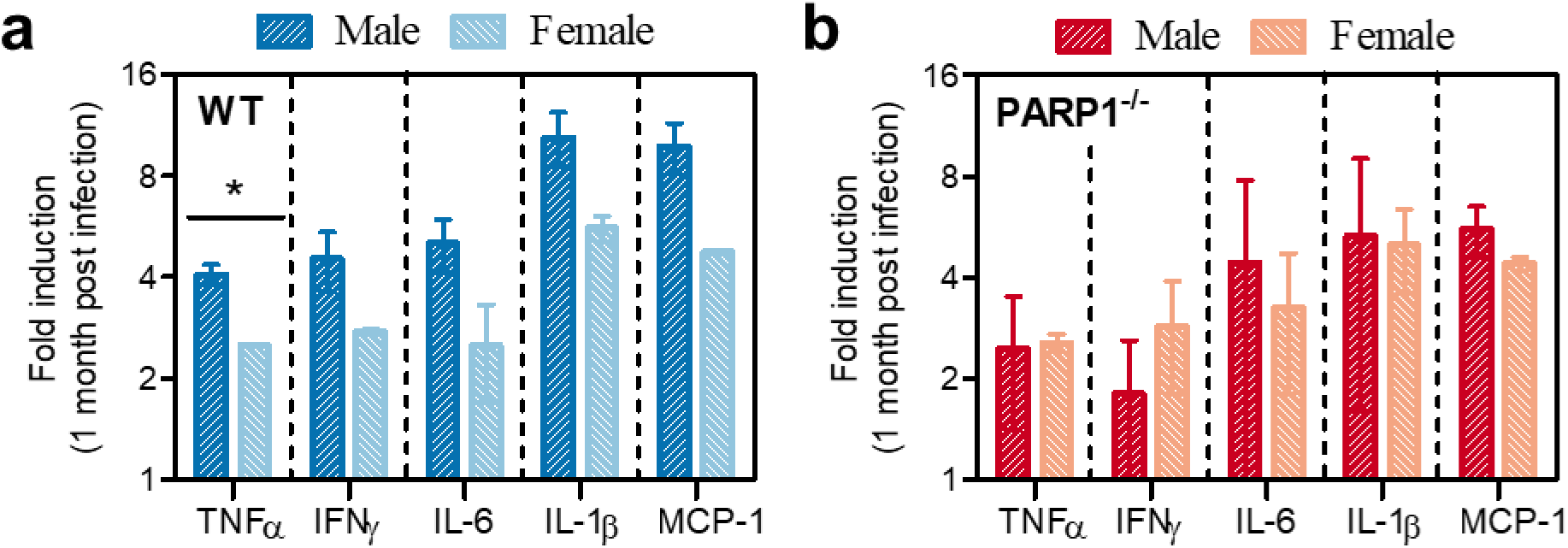
PARP1 deletion abolishes sex differences in TB immune responses. Induction of key cytokines and chemokines in the lungs of male and female (**a**) WT (blue) and (**b**) PARP1^-/-^ (red) mice 28 days after aerosol infection with *M. tb* H37Rv (implantation: 1.86 log_10_ CFU). Lung cyto-/chemokines were quantified by Luminex multiplex assay and are expressed as fold change relative to levels in uninfected lungs (n=2-3). Values above 1 indicate levels that are higher in infected than in uninfected lungs. *, *p* < 0.05 by unpaired t-test. PARP1 deletion reduced the induction of TNFα, IFNγ, IL-6, IL-1β and MCP-1 in male mice to female levels and eliminated the sex differences observed in WT mice.

Male WT mice displayed enhanced proinflammatory TB responses compared to PARP1^-/-^ males early in infection (28 dpi) but not in the chronic phase (74 dpi) (**Fig. 4a; Supplementary Fig. S3c-d**). Interestingly, PARP1 deletion only impaired proinflammatory responses in male but not in female mice (**Fig. 4a-b**), suggesting that PARP1 enhances the production of proinflammatory mediators in TB in a sex- and time-dependent manner. In contrast, induction of IL-10, a regulatory cytokine that is modestly induced in TB and associated with improved containment and reduced tissue damage, was absent in both male and female PARP1^-/-^ mice, indicating that PARP1 is required for IL-10 production in a sex-independent manner (**Fig. 4c-d**) (42). Together, these data show that PARP1 deletion dampened early proinflammatory immune responses, including TNFα, IL-1ß and IFNγ production specifically in male mice, but eliminated IL-10 responses in both male and female mice.

**Figure 4.**
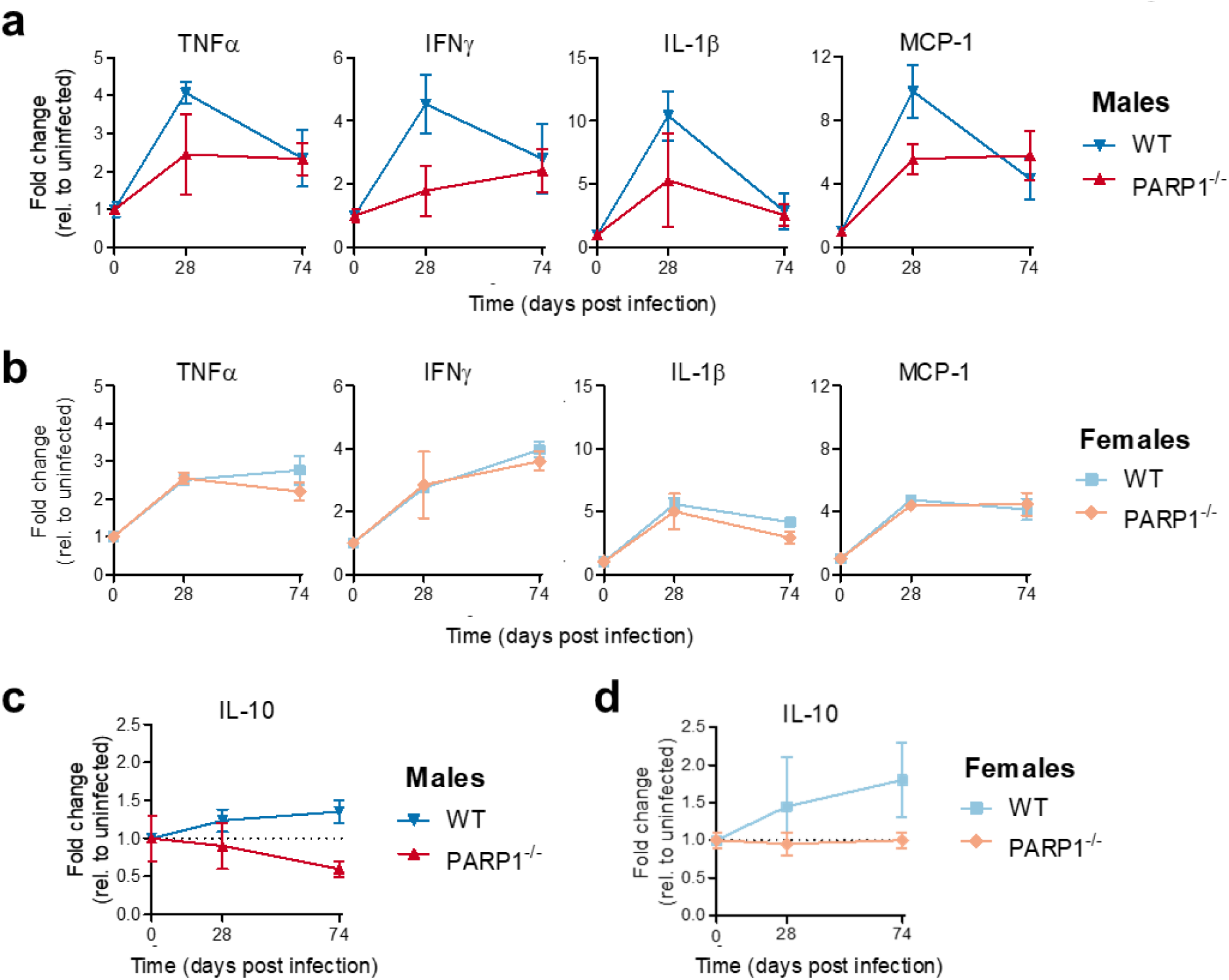
PARP1 deletion dampens proinflammatory responses in male but not in female mice. Change in lung cytokines and chemokines over time in (**a,c**) male or (**b,d**) female WT (blue) or PARP1^-/-^ (red) mice following aerosol infection with *M. tb* H37Rv (implantation: 1.86 log_10_ CFU). Lung cytokines were quantified by Luminex multiplex assay and are expressed as fold change relative to levels in uninfected lungs (n=2-3). Values above 1 indicate levels that are higher in infected than in uninfected lungs. PARP1 deletion dampened IL-1ß, TNFα, IFNγ and MCP-1 responses following *M. tb* infection in males but not in females but eliminated IL-10 induction in both male and female mice.

### PARP1 deletion is protective and improves TB containment in female mice

Since PARP1 enhances proinflammatory responses important for TB containment in male but not in female mice, we hypothesized that PARP1^-/-^ mice, in particular males, would be more susceptible to TB than WT mice. We therefore compared survival and disease progression in WT and PARP1^-/-^ mice following aerosol infection with *M. tb* H37Rv. Unexpectedly, PARP1^-/-^ mice appeared to be somewhat protected and survived moderate infection longer than WT mice (median survival: 109.5 in PARP1^-/-^ vs. 93 days in WT mice; not significant) (**Table 1; Fig. 5**). Surprisingly, we discovered that this protective effect of PARP1 deletion was due to the profound resistance of PARP1^-/-^ females, who survived infection significantly longer than WT females (134 vs. 83.5 days, respectively; *p*=0.0031; hazard ratio 5.225) (**Fig. 5b**; **Table 1**). In contrast, survival did not differ between WT and PARP1^-/-^ males (**Fig. 5c; Table 1**). These data indicate that PARP1 deletion has sexually divergent effects and prolongs female survival in TB.

**Figure 5.**
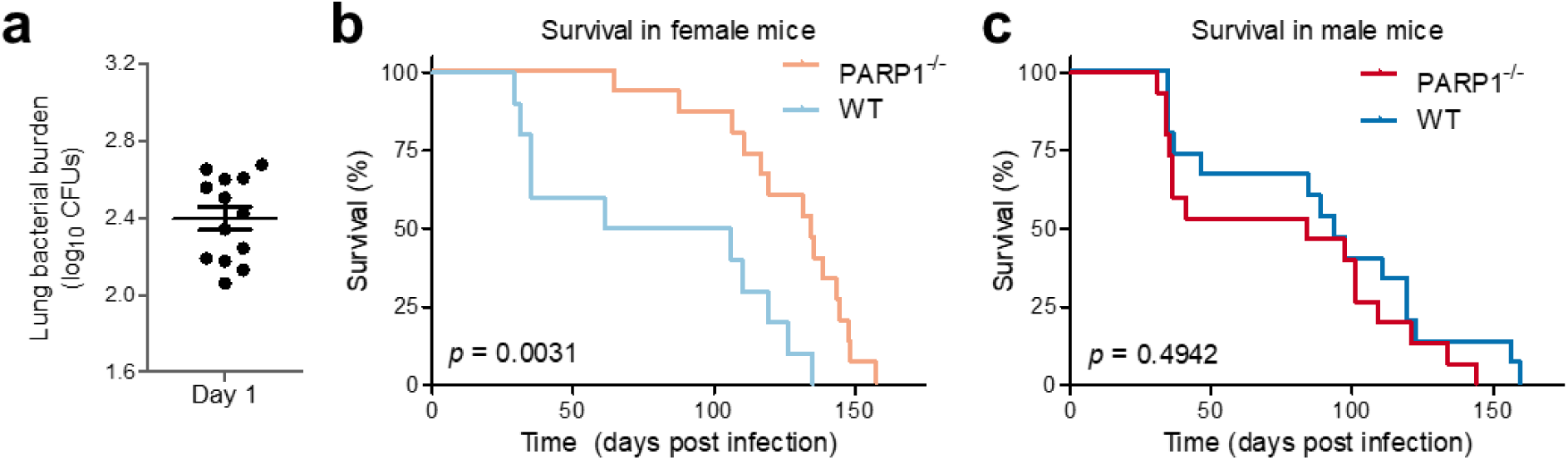
PARP1 deletion prolongs TB survival in female mice. TB survival in WT (blue) or PARP1^-/-^ (red) mice aerosol-infected with *M. tb* H37Rv. (a) Day 1 lung implantation (mean: 2.397 log_10_ CFUs). (b-c) Survival analysis in *M. tb*-infected female (**b**; n=10-15) or male (**c**; n=15) WT or PARP1^-/-^ mice Statistical differences in survival were assessed by Gehan-Breslow-Wilcoxon test. PARP1^-/-^ females survive infection significantly longer than WT females.

**Table 1:** Survival statistics follwing moderate TB infection (2.397 log_10_ CFU).

| **Table 1:** Survival statistics following moderate TB infection (2.397 log<sub>10</sub> CFU). |  |  |  |  |
| --- | --- | --- | --- | --- |
|  | **Median Survival (days)** |  | **Hazard Ratio**<br>(95% CI) | **Significant?**<br>(*p*) |
|  | **WT** | **PARP1<sup>-/-</sup>** |  |  |
| **Females** | 83.5 | 134 | 5.225<br>(1.758 – 15.52) | **Yes**<br>**(0.0031)** |
| **Males** | 93 | 84 | 0.7030<br>(0.3248 – 1.522) | No<br>(0.4942) |
| **Combined** | 93 | 109.5 | 1.325<br>(0.7497 – 2.342) | No<br>(0.1375) |

In line with these observations, PARP1^-/-^ females contained *M. tb.* proliferation more efficiently than PARP^-/-^ males and never exceeded a lung bacterial burden of 6.7 ± 0.3 log_10_ CFUs (**Fig. 6a**, left). By comparison, the lung bacterial burdens peaked at 7.3 ± 0.3 log_10_ CFUs in WT females (28 dpi), at 7.2 ± 0.3 log_10_ CFUs in WT males (42 dpi) and at 7.3 ± 0.5 log_10_ CFUs in PARP1^-/-^ males (28 dpi). Similarly, the bacterial burden in the spleen tended to be lowest in PARP1^-/-^ females and highest in PARP1^-/-^ males (**Fig. 6a**, right). We consistently observed these same patterns in three independent experiments with similar infection parameters and day 1 lung implantations ranging from 1.66-1.86 log_10_ CFUs. The combined analysis of these studies confirmed that PARP1^-/-^ females have significantly fewer lung bacteria than WT females one month after infection (*p*<0.05), resulting in an overall lower bacterial burden in PARP1^-/-^ mice (**Fig. 6b**). These results suggest that PARP1 deletion is beneficial in female mice and increases TB survival by enhancing the control of bacterial replication.

**Figure 6.**
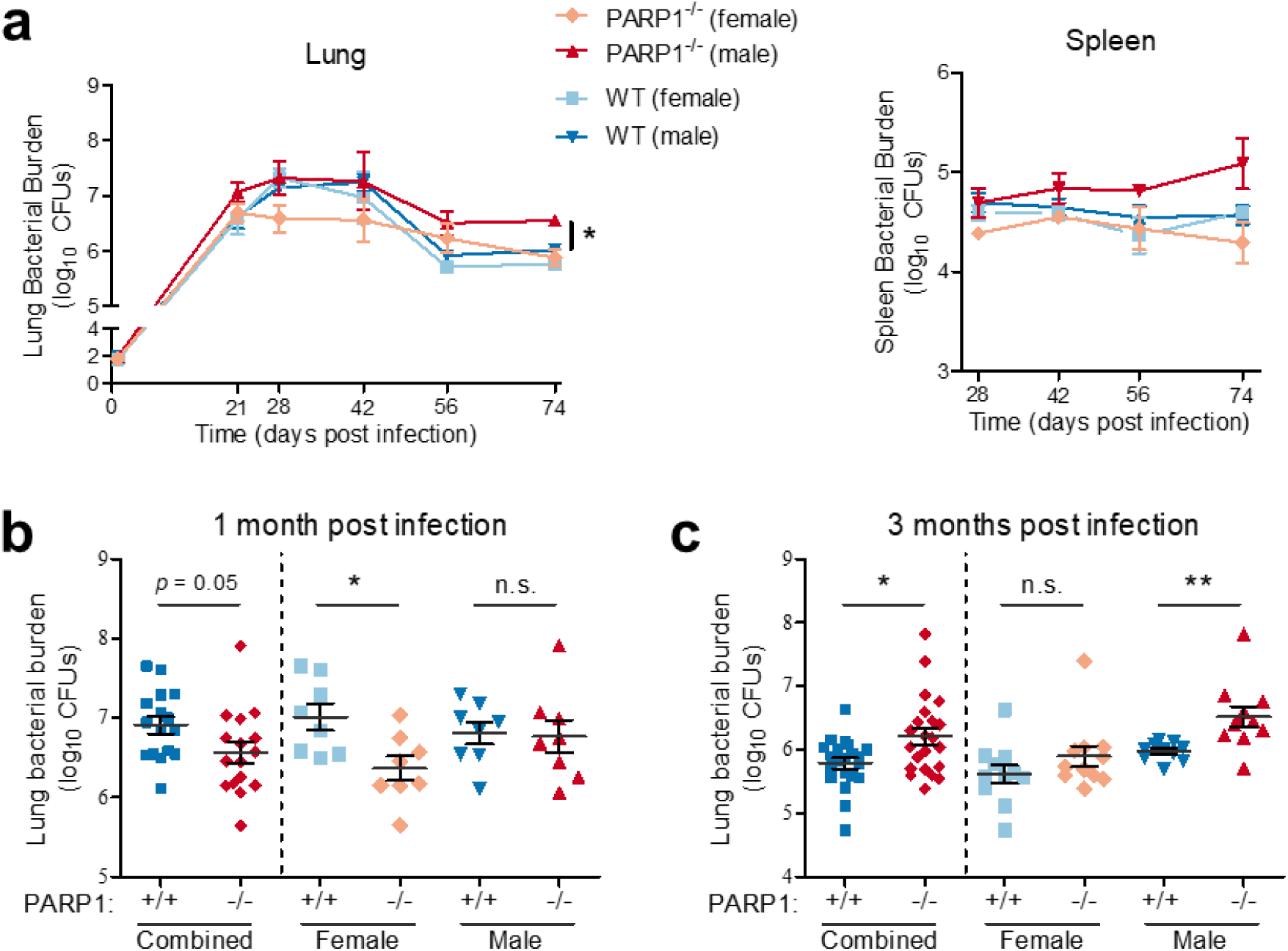
Bacterial containment is improved in female but impaired in male PARP1-deficient mice. Bacterial burden in WT (blue) or PARP1^-/-^ (red) mice aerosol-infected with *M. tb* H37Rv. (a) Lung (left) and spleen (right) bacterial burden in female or male WT or PARP1^-/-^ mice (day 1: 1.86 log_10_ CFU). Each point represents the mean bacterial burden of 2-3 male or female mice. *, *p* < 0.05 by unpaired t-test (lung CFUs: male vs. female PARP1^-/-^ mice). (b-c) Lung bacterial burden in WT (blue; +/+) and PARP1^-/-^ (red; -/-) mice after (b) 1 or (c) 3 months of infection with *M. tb* H37Rv or H37RvΔ*pncA.* Data was collected in three independent experiments (day 1: 1.659 (0.3172), 1.724 (0.0916) and 1.858 (0.2356) log_10_ CFU (SD)). Horizontal lines represent the mean +/-SEM. *, *p* < 0.05; **, *p* < 0.01 by unpaired t-test. PARP1 deficiency improves bacterial containment in females but impairs it in male mice.

### PARP1 deletion impairs TB control and exacerbates inflammation in male mice

Unlike females, PARP1^-/-^ males contained early *M. tb.* infection similar to WT mice but developed higher lung and spleen bacterial burdens than any other group after 2.5 – 3 months of infection (**Fig. 6**). As a result, PARP1^-/-^ mice (males and females combined) had significantly more lung bacteria than WT mice by three months post infection (*p*<0.05), driven by the high bacterial burden in PARP1^-/-^ males (*p*<0.01) (**Fig. 6c**). Even though TB survival did not differ significantly in males, the hazard ratios indicate that the risk of death was nearly 30% lower in WT males than in PARP1^-/-^ males following infection (**Table 1**). Therefore, in contrast to females, PARP1 deletion is detrimental in males and antagonizes bacterial containment.

While there were no striking differences in gross pathology, the lungs of PARP1^-/-^ males appeared notably enlarged after 2-3 months of infection in all experiments (**Fig. 7a**; **Supplementary Fig. S4**). In fact, after three months of infection, the lungs of PARP1^-/-^ males were approximately 1.5-times heavier than those of any other group, even though PARP1^-/-^ males weighed less than WT males and the lung weights of age-matched uninfected mice were comparable across all groups (**Fig. 7b-c**). Histopathologic analysis revealed that PARP1^-/-^ males had significantly more lung inflammation and consolidation than mice of any other group (*p*<0.05) (**Fig. 7d-f**; **Supplementary Fig. S5**). We also assessed lung inflammation in 3 additional mice by ^18^F-FDG PET and by CT but did not observe significant differences between groups (**Supplementary Fig. S6**). Together, these data suggest that PARP1 deletion impairs immune clearance of *M. tb* and increases lung inflammation during chronic TB in males but not in females.

**Figure 7.**
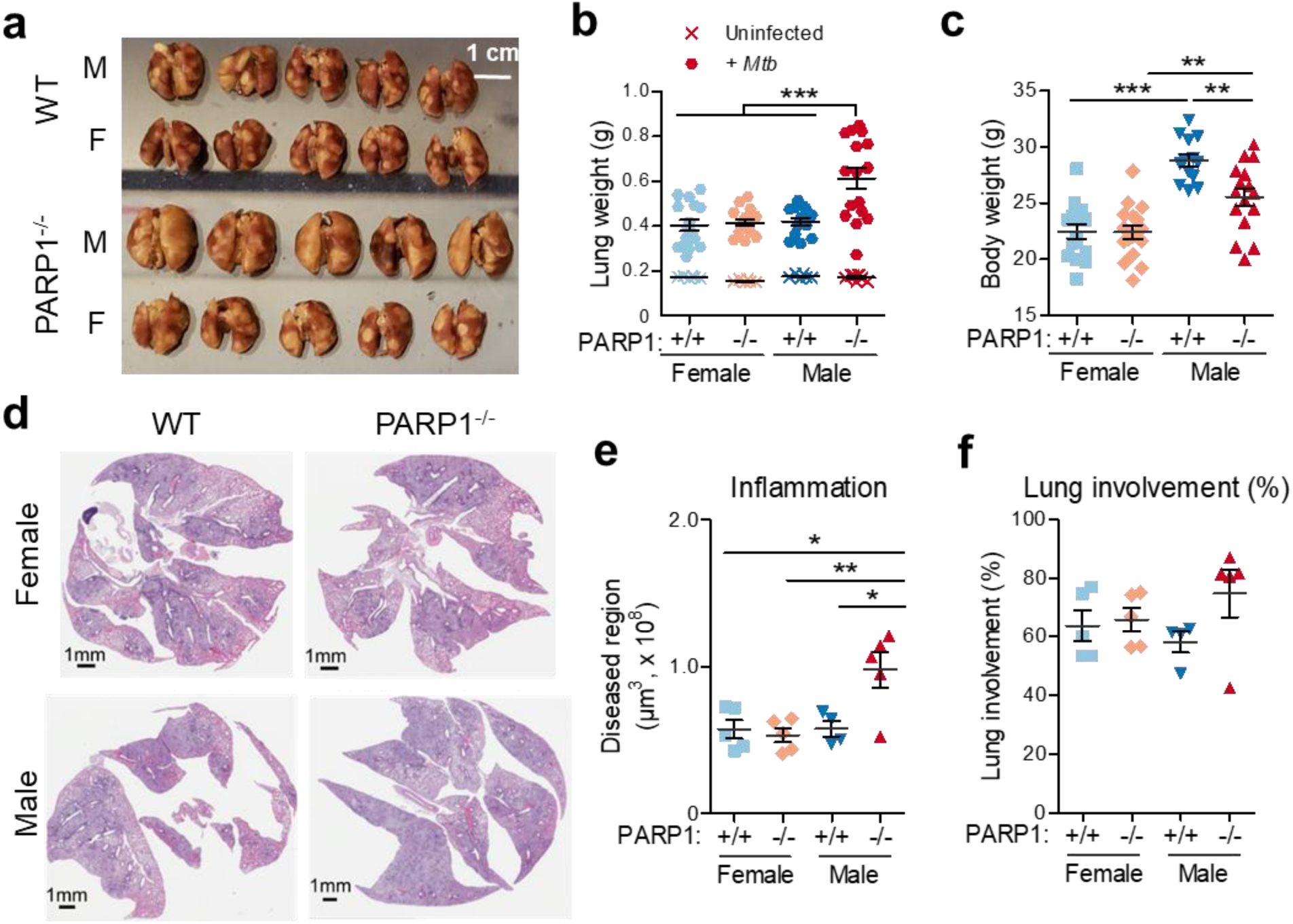
PARP1 deletion exacerbates TB lung inflammation in male mice. (a) Formalin-fixed lungs of male (M) and female (F) WT or PARP1^-/-^ mice 3 months after infection with *M. tb* H37Rv (lung implantation: 1.66 log_10_ CFU). (b) Fresh (unfixed) lung weights of male and female WT (blue) or PARP1^-/-^ (red) mice 3 months after *M. tb* infection (circle) or of age-matched uninfected controls (cross). (c) Corresponding body weights of the infected mice depicted in (b). *, *p* < 0.05; ***, *p* < 0.001 by student’s t-test. The lungs of infected, but not uninfected, PARP1^-/-^ males are significantly larger than those of any other group, despite them having a lower total body weight than WT males. (d) Representative H&E-stained lung sections of male and female WT or PARP1^-/-^ mice 3 months after infection with *M. tb* H37Rv (lung implantation: 1.66 log_10_ CFU). There was extensive consolidation throughout the lungs, in particular in PARP1^-/-^ males. (e-f) Quantification of area of consolidation (e) and percent lung involvement (f) indicating lung inflammation. Each point represents an individual mouse. One-way analysis of variance with Bonferroni post-test; *, *p* < 0.05; **, *p* < 0.01. PARP1^-/-^ males have significantly more lung inflammation than females or WT males in chronic TB.

### PARP1 contributes to TB sex differences by driving sexually divergent immune responses

Consistent with published results, WT males had enhanced proinflammatory TB responses early in infection (28 dpi) but similar responses to females in the chronic phase (74 dpi) (**Fig. 3a**; **Supplementary Fig. S3c**) (9). Despite these sex differences in immune responses, TB progression and inflammation did not differ between male and female 129S1 WT mice (**Supplementary Fig. S7**), suggesting that there might be fundamental differences in the mechanism of TB containment in males and in females. PARP1 deletion eliminated the sex difference in immune responses (**Fig. 3b**) but impaired TB containment in males and improved it in females (**Fig. 2a; Fig. 8**). Therefore, the PARP1-dependent enhancement of early male cytokine responses may be a compensatory mechanism required to achieve comparable TB containment in males and females.

**Figure 8.**
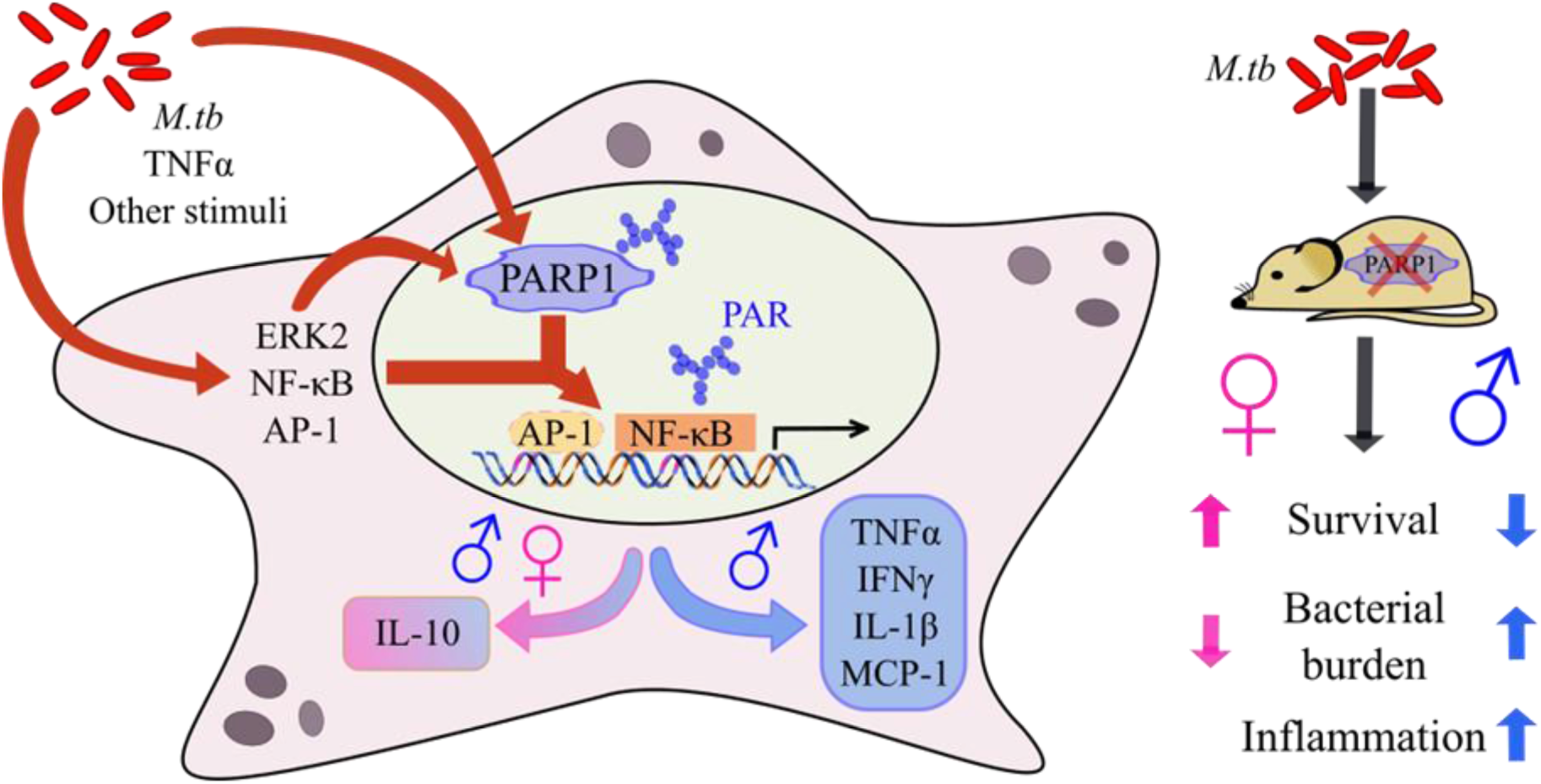
Proposed model: PARP1 drives TB sex differences. PARP1 is activated by *M. tb* infection and interacts with numerous transcription factors, such as ERK2, NF-κB and AP-1, to shape the TB host response. PARP1 drives exaggerated inflammatory responses in males that enhance their ability to contain the infection. In contrast, PARP1 antagonizes TB containment in females by promoting primarily immunosuppressive IL-10 responses. As a result, PARP1 deletion increases TB resistance in female mice but exacerbates inflammation in male mice.

## Discussion

TB has a male bias, associated primarily with biological rather than behavioral factors, but the underlying mechanisms are largely unknown (4). Here, we identified the master regulator PARP1 as a nuclear factor that contributes to sexually divergent TB immune responses and disease susceptibility. Despite comparable rates of infection, men progress to active TB not only more frequently but also with higher severity and mortality than women (3–7). A male bias in TB severity and mortality has also been reported in C57BL/6 and BALB/c mice; curiously, this was associated with increased early proinflammatory immune responses in male C57BL/6 mice whereas in BALB/c mice, the inverse was observed (9, 10). While we saw no clear sex differences in TB progression in 129S1/SvlmJ mice, as has been described in C57BL/6 mice, we observed markedly enhanced production of proinflammatory mediators after one month in infected males that depended on the presence of functional PARP1. Interestingly, a similar role for sexually divergent immune regulation has been described for PARP1 in a mouse model of LPS-induced systemic inflammation: while females are highly resistant to LPS, males produce high levels of TNFα and die rapidly after LPS exposure, but PARP1 deletion or inhibition reduces TNFα levels and death in males to female levels (29). Therefore, PARP1 might also drive TB sex differences via the differential regulation of male and female immune responses, particularly during early infection.

Since high levels of inflammatory cytokines can contribute to bacterial containment but also promote immunopathology, exaggerated immune responses might thus be associated with worse TB outcomes. However, our findings indicate that robust proinflammatory responses early in infection are critical for long-term TB containment in males but not in females. Lack of PARP1 in male mice lowered cytokine responses to female levels but was associated with impaired bacterial control and paradoxically exacerbated inflammation during the chronic phase of infection. Since PARP1 drives proinflammatory responses in males, we expected WT males to have higher levels of tissue damage and lung inflammation and were surprised to observe more inflammation in PARP1^-/-^ males despite reduced proinflammatory responses. One explanation for these observations is that TB control in males might rely more heavily on cytokine-driven early bactericidal mechanisms than in females. Attenuation of this capacity in PARP1^-/-^ males might limit their ability to restrict bacterial replication, over time resulting in increased dissemination and lung damage. Therefore, males might compensate for fundamental sex differences in immune functions that are disadvantageous in TB by boosting early bactericidal mechanisms with the PARP1-dependent enhancement of proinflammatory mediators (**Fig. 8**).

Another explanation for the increased bacterial burden and inflammation in PARP1^-/-^ males is the increased production of MCP-1 during the chronic phase of infection, by which time most immune responses were similar in all groups (**Fig. 4; Supplementary Fig. S3; Supplementary Fig. S8**). In humans, high MCP-1 levels are associated with an increased risk of TB progression, and we observed a tight correlation between MCP-1 levels and bacterial burden (**Supplementary Fig. S8**) (43). MCP-1 might play dual roles in the control and expansion of mycobacterial infection and inadvertently benefit the pathogen by increasing the pool of infectable macrophages. Since MCP-1 induction was consistently low in PARP1^-/-^ females, the protective effects of PARP1 deletion in females might similarly be a result of reduced macrophage migration to the lungs.

Alternatively, the increased inflammation in PARP1^-/-^ males might be a result of their absent IL-10 induction. IL-10 is a regulatory cytokine that is modestly induced during TB and antagonizes the proinflammatory effects of TNFα, IFNγ, and IL-12p40 (42). However, proinflammatory responses in PARP1^-/-^ males were reduced rather than increased despite the absence of IL-10, indicating that PARP1 likely regulates anti- and proinflammatory immune responses via independent mechanisms. Even though IL-10 induction was PARP1-dependent in males and in females, the absence of IL-10 might impact TB progression differently in males and in females depending on the respective immune context. IL-10 can benefit both the pathogen, *e.g.,* by suppressing innate and adaptive immune responses, and the host, *e.g.,* by reducing tissue damage (44). In contrast to male mice, PARP1 deletion was profoundly protective in females, who exhibited significantly lower bacterial counts and prolonged survival than their WT counterparts. The only observed difference in immune responses between WT and PARP1^-/-^ females was the lack of IL-10 induction in PARP1^-/-^ mice (**Fig. 4**). Since IL-10 deletion or inhibition is associated with increased bacterial restriction and survival in mouse models of TB, lack of IL-10 could account for the improved survival and bacterial containment in PARP1^-/-^ females (42). The bacterial burden was most reduced In PARP1^-/-^ females one month into infection, around the time when the initial period of unrestricted bacterial replication ends due to the onset of adaptive immunity (45). Therefore, the TB resistance of PARP1^-/-^ females might be the result of improved activation of adaptive immune responses, potentially because of absent immunosuppressive effects of IL-10. Importantly, since PARP1 deletion was profoundly protective in infected females, adjunctive PARP inhibition has the potential to improve TB clearance, particularly in women.

While our study revealed new insight into TB sex differences, further underlying mechanisms behind the male bias remain to be elucidated. The focus of our study was the comparison of disease progression and immune responses in WT and PARP1^-/-^ mice. However, it is entirely possible that the sexually dimorphic phenotype of PARP1 deletion in TB is only indirectly related to its function as an immune regulator. For example, PARP1 might drive TB sex differences via sexually dimorphic cell death paradigms. PARP1 regulates cell death signaling and acts as a switch between cell fates: PARP1 inhibits caspase activation and is reciprocally inactivated by caspases; PARP1 can induce caspase-independent, apoptotic as well as various forms of necrotic cell death; and PARP1 is implicated in RIPK-dependent and -independent forms of necroptosis (38, 46). In TB, apoptosis is host-protective by killing intracellular bacteria and stimulating adaptive immunity while necrosis is associated with bacterial dissemination and disease severity (44). Interestingly, cell death is sexually dimorphic and predominantly caspase-mediated in females but caspase-independent, PARP1-mediated in males (27, 31, 34, 47). Accordingly, PARP1 deletion reduces cell death in males but increases caspase activity and cell death in females (27, 34, 47). Importantly, PARP inhibition reduces necrotic and increases apoptotic and live cell population, thereby shifting from a necrotic (unfavorable in TB) to an apoptotic (host-protective) milieu (28). Therefore, TB sex differences might be the result of divergent cell death modalities that were highlighted by PARP1 deletion. For example, by enhancing apoptosis in *M. tb*-infected females, PARP1 deficiency could potentially promote adaptive immune activation and accelerate TB containment, resulting in lower bacterial burden in female mice around one month post infection that remain well controlled throughout the chronic phase. Additional studies are needed to determine whether divergent cell death paradigms contribute to TB sex differences in a PARP1-dependent or -independent manner. In addition, our study did not assess the hormonal contribution toward the male bias in TB. PARP1 is indispensable for steroid sex hormone signaling, and sex hormones reciprocally activate PARP1 (32, 33, 38, 48). TB infection studies utilizing castrated and ovariectomized mice and sex hormone supplementation might therefore yield additional insight into the role of sex hormones to TB sex differences.

In summary, we showed that PARP1 is robustly activated during *M. tb.* infection and contributes to immune responses and the control of infection in a sexually divergent manner. In contrast to LPS-induced inflammation or infarct models, where PARP1 deficiency mainly benefits males, PARP1 deletion in TB was protective in female but detrimental in male mice (28, 29). The impact of PARP1 deficiency in TB was also less pronounced than in other proinflammatory conditions, such as endotoxic shock and LPS-induced lung injury (17, 49). These contrasting results likely reflect the disproportionate complexity of host-pathogen interactions over the course of TB infection, not all of which are recapitulated in a mouse model. The fact that PARP1 activity in healthy human volunteers is significantly higher in men than in women supports a sexually divergent role of PARP1 in humans (28, 32). Based on our findings, we propose that in TB PARP1 is host-protective in males by enhancing early pro-inflammatory cytokine responses, but antagonistic in females which except for IL-10 do not show major changes in PARP1-dependent cytokine expression. Since PARP1 deletion was profoundly protective in infected female mice, adjunctive PARP inhibition may have the potential to improve TB clearance, particularly in women. In conclusion, PARP1 deletion revealed fundamental differences in the male and female response to TB and might pave the way to a better understanding of TB sex differences.

## Methods

### Animals

Six-week-old female C57BL/6J (stock #000664) and C3HeB/FeJ (stock #658) mice were purchased from The Jackson Laboratory, and female BALB/c (strain #028) and 129S2/SvPasCrl (strain #476) mice from Charles River Laboratories. PARP1-deficient 129S-*Parp1^tm1Zqw^*/J mice (stock #002779, Jackson Labs) were bred in-house. PARP1 disruption was routinely validated by PCR as described by the supplier (protocol 22839, version 2.3) using the common forward primer (5’-CATGTTCGATGGGAAAGTCCC-3’), a WT reverse primer (5’-CCAGCGCAGCTCAGAGAAGCCA-3’) and a mutant reverse primer (5’-AGGTGAGATGACAGGAGATC-3’). Age-matched male and female recommended control 129S1/SvlmJ mice (stock #002448, Jackson Labs) were purchased for each experiment. All animal procedures were approved by the Institutional Animal Care and Use Committee of the Johns Hopkins University School of Medicine.

### Bacterial strains

*M. tuberculosis* strains H37Rv and H37RvΔ*pncA* were obtained from the Johns Hopkins Center for Tuberculosis Research, grown to an optical density at 600 nm of approximately 1.0 in Middlebrook 7H9 broth (Gibco) supplemented with 10% (v/v) oleic acid-albumin-dextrose-catalase (OADC; Difco), 0.5% (v/v) glycerol and 0.05% (v/v) Tween 80 (Sigma-Aldrich) and stored in 1 ml aliquots at −0°C.

### Aerosol infections

Mice were infected between 8 and 12 weeks of age via the aerosol route using the Glas-Col Inhalation Exposure System (Terre Haute, IN). A new stock of *M. tuberculosis* was used for each infection and diluted in sterile phosphate-buffered saline (PBS, pH 7.4) or 7H9 broth with OADC, glycerol, and Tween 80 at empirically determined factors to achieve the desired inoculum. For high-dose infections (day 1 implantation >2.5 log_10_ CFUs), mycobacteria were thawed one week prior to infection and sub-cultured to achieve mid-log phase on the day of infection. For low-dose infections (day 1 implantation <2.5 log_10_ CFUs), a mycobacterial stock was thawed on the day of infection and used immediately. To reduce intergroup variability, mice from all comparative groups were infected together or evenly distributed between infection cycles (*i.e.*, equal numbers of male and female PARP1^-/-^ and WT mice were infected in the same cycle). On the day after infection, 3-5 mice per cycle were sacrificed to determine the number of CFUs implanted into the lungs. The general appearance and body weight of mice were monitored at least weekly throughout all experiments. All infections, housing of infected mice and handling of infectious materials were carried out under biosafety level 3 containment in dedicated facilities.

### PARP1 inhibition in mice

Talazoparib (BMN-673; CAS no. 1207456-01-6, Medchemexpress LLC) was reconstituted in HPLC-grade DMSO (Sigma-Aldrich) and stored in 10 mM aliquots at −20°C. DMSO aliquots were frozen at the same time to prepare vehicle control solutions. For PARP inhibition in mice, a talazoparib solution was prepared fresh daily in 0.5% low-viscosity carboxylmethyl cellulose (Sigma-Aldrich) and administered by orogastric gavage once-daily for 5 consecutive days, in a total volume of 0.2 ml per treatment.

### Tissue collection and bacterial enumeration

Mice were sacrificed at predetermined intervals and the total body weight was recorded. Lungs and spleens were aseptically removed and sectioned for bacterial enumeration (A; right lung lobes) and PAR, cytokine and chemokine quantification (B; left lung lobes). The weights of the intact lung/spleen and the (B) section were recorded and used to estimate total bacterial burden as described below. For bacterial enumeration, lungs/spleens (A) were placed in 2.5 ml sterile PBS for 24-48 h at 4°C, examined for gross pathology and manually homogenized. Homogenates were serial-diluted, and 0.5 ml plated on Middlebrook 7H11 agar (Difco) supplemented with 10% (v/v) OADC, 0.5% (v/v) glycerol, 10 mg/ml cycloheximide, 50 mg/ml carbenicillin, 25 mg/ml polymyxin B and 20 mg/ml trimethoprim (Sigma-Aldrich). Plates were incubated at 37°C for 3-4 weeks before colonies were counted. Colony numbers were adjusted by the plating, dilution and dissection factors and log-transformed to estimate the total colony-forming units (CFUs) per organ (CFUs = log_10_ [(number of colonies)*(5)*(A+B) / (A)] + [dilution factor]).

### Immunoblot and PAR analysis

PARP1 activity was assessed by comparing poly(ADP-ribose) (PAR) levels in experimental and control samples by immunoblot (24, 39). Lung sections were placed in chilled extraction buffer (50 mM glucose, 25 mM Tris-HCl pH 8.0, 10 mM EDTA) with protease inhibitor cocktail (Sigma-Aldrich) and 1.0 mm zirconia beads (5.5 g/cc; Biospec Products) and immediately homogenized by bead-beating in a mini-beadbeater (Biospec Products) in 30 sec intervals at 4,800 RPM. Homogenates were placed on ice for 10 min, mixed with denaturing sample buffer (69.45 mM Tris-HCl pH 6.8, 11.1% (v/v) glycerol, 1.1% (v/v) LDS, 0.005% (v/v) bromophenol blue, 2.5% (v/v) ß-mercaptoethanol) and denatured at 90°C for 15 min. Lysates were chilled on ice and stored at −80°C for immunoblot analysis.

Protein concentrations were determined by CB-X protein assay (G-Biosciences) according to the manufacturer’s instructions. Cell lysates or lung homogenates in denaturing sample buffer (10-20 µg total protein) were separated by SDS-PAGE on 4-15% Mini Protean TGX gels (Bio-Rad) and transferred electrophoretically to a 0.45 µm nitrocellulose blotting membrane (GE Healthcare) at 100 V for 75 min at 4°C. To assess transfer efficiency, membrane was immersed in 0.1% (w/v) Ponceau S in 5% (v/v) acetic acid for 5 min, rinsed with distilled water and imaged. Membrane was de-stained in 0.1 M sodium hydroxide, rinsed with running distilled water for 2-3 min and washed with tris-buffered saline containing 0.1% (v/v) Tween 20 (TBST). The membrane was then blocked in TBST containing 5% (w/v) nonfat dry milk for 1 h at room temperature (RT), washed in TBST and incubated with a recombinant human monoclonal anti-PAR antibody (1:2,500 in 5% nonfat dry milk-TBST; clones #19 and #21, highly specific for target-bound PAR, custom-designed at Bio-Rad AbD Serotec GmbH) (39) with gentle agitation overnight at 4°C. Following primary antibody incubation, the membrane was washed, incubated with HRP-conjugated goat anti-human IgG (Fab’)2 (1:5,000 in 1% nonfat dry milk-TBST; Abcam) for 1 h at RT and visualized with KwikQuant Ultra digital ECL substrate using the KwikQuant digital imager (Kindle Biosciences, LLC).

After visualizing PAR bands, the membrane was washed with TBS and stripped in a 200 mM glycine solution (pH 2.5) for 1 h at RT. The membrane was washed with TBS and TBST, blocked in TBST containing 5% (w/v) nonfat dry milk for 1 h at RT, washed with TBST, incubated with HRP-conjugated mouse monoclonal anti-beta Actin antibody [AC-15] (1:50,000 in 5% (w/v) BSA in TBST; Abcam) for 0.5-1 h at RT and visualized as described for PAR detection. For PARP1 protein detection, membrane was incubated with the primary HRP-conjugated (1:1,000 in 5% nonfat dry milk; Cell Signaling Technology) overnight at 4°C and secondary HRP-conjugated anti-rabbit IgG (1:2,000 in 5% nonfat dry milk; Cell Signaling Technology) for 1 h at RT. Digital images were converted to black-and-white using Photoshop, and relative band intensities were quantified using ImageJ (version 1.52a) as described (50). PARP1 activity was defined as the intensity of high-molecular weight (72 – 250 kDa) PAR bands after normalizing to β-actin and was expressed as fold change relative to uninfected or untreated control samples on the same immunoblot.

### Cytokine and chemokine analysis

Lung cytokines/chemokines (GMCSF, IFNγ, IL-1ß, IL-2, IL-6, IL-10, IL-12(p70), IL-17/CTLA8, MCP-1/CCL2, MIP-1α/CCL3, TNFα) were analyzed by Luminex multiplex bead assay on a Bio-Plex 200 platform (Bio-Rad) with a mouse cytokine/chemokine magnetic bead panel (MCYTOMAG-70K, lot # 3224392; Millipore) according to the manufacturer’s instructions. Lung tissue was disaggregated in a mini-beadbeater (Biospec Products) at 4,800 RPM with 1.0 mm zirconia beads (5.5 g/cc; Biospec Products) in sterile PBS, incubated on ice for 5 min and centrifuged at 12,000 x*g* for 10 min. Supernatants were filter-sterilized through 0.22 µm cellulose acetate Spin-X centrifuge tube filters (Costar) and stored at −80°C. Protein concentrations were determined by CB-X protein assay (G-Biosciences) according to the manufacturer’s instructions, and samples were diluted to 1 mg/ml for analysis. Results are presented as concentrations (pg/ml) or fold change relative to uninfected lungs from the same group.

### Histopathology and inflammation analysis

For histology, intact lungs were fixed by immersion in 10% neutral-buffered formalin for 48 hours, paraffin-embedded, sectioned and stained with hematoxylin and eosin (H&E), Masson’s trichrome or Ziehl-Neelsen stain for acid-fast bacilli. Slides were digitally scanned at 40x on an Aperio AT turbo scanner console version 102.0.7.5 (Leica Biosystems). Image files were transferred using Concentriq for Research version 2.2.4 (Proscia Inc), randomized and scored blinded to experimental grouping using Aperio ImageScope (Leica Biosystems). Regions of interest (ROI) were manually selected at 20x magnification. Inflammatory regions were defined by leukocyte disruption of alveolar architecture. Lung involvement was calculated as a percentage of the total lung area.

For ^18^F-FDG PET/CT, live animals were imaged at 84 days post-infection inside in-house developed, sealed biocontainment devices compliant with BSL-3 isolation (51). Mice were fasted overnight, and ^18^F-FDG (5.31 ± 0.25 MBq) was administered intravenously via the tail vein. A 15-minute PET acquisition and CT were performed using the nanoScan PET/CT (Mediso, Washington, DC) 45 min after tracer injection. Volumes of interest (VOIs) were manually selected using CT as a guide and applied to the PET dataset using VivoQuant 2020 (Invicro, Boston, MA). Mean lung ^18^F-FDG PET activity was calculated for each mouse as the average activity of all VOIs normalized by injected dose and body weight and is expressed as standard uptake values (SUV).

### Statistical analysis

Statistical analyses were performed using Prism version 5.01 for Windows (GraphPad) or Microsoft Excel (Microsoft Office Professional Plus 2019). Statistical tests used are indicated in the figure legends. Differences between two group means were assessed by unpaired, two-tailed *t*-test, with Welch’s correction for unequal variance where indicated. Data sets containing three or more groups were analyzed by one-way analysis of variance with Bonferroni post-hoc test. Survival curves were compared by Gehan-Breslow-Wilcoxon test. CFU counts were log_10_-transformed prior to analysis. A *p* value below 0.05 was considered significant. Data represent mean ± SEM.

### Data availability

All data generated during the current study are available from the corresponding author on reasonable request.

## Acknowledgements

This work was supported by the JPB foundation and National Institute of Health (NIH) grants AI 137659, AI 37856, AI 130595, HL 133190, HL 140812, P50 NS 38377 and 1RF1 AG 059686. T.M.D. is the Leonard and Madlyn Abramson Professor in Neurodegenerative Diseases. We thank Elizabeth Ihms, DVM, PhD, DACVP (Dane & Dutchy LLC) and Chris Thoburn (SKCCC Immune Monitoring Core) for helpful discussions and assistance with histopathology and for cytokine/chemokine analysis, respectively.

## Author Contributions

W.R.B, V.L.D. and S.K. conceived the project. W.R.B., S.K., V.L.D., B.K., S.K.J. and A.A.O. designed the experiments. S.K., A.A.O. and M.K. carried out experiments. T.M.D., V.L.D. and B.K. provided critical reagents and technical expertise. S.K. analyzed the data and prepared the tables and figures. W.R.B and S.K. interpreted the data and wrote the manuscript. All authors read, provided feedback on, and approved the manuscript.

## Supplemental Figures

**Figure S1.**
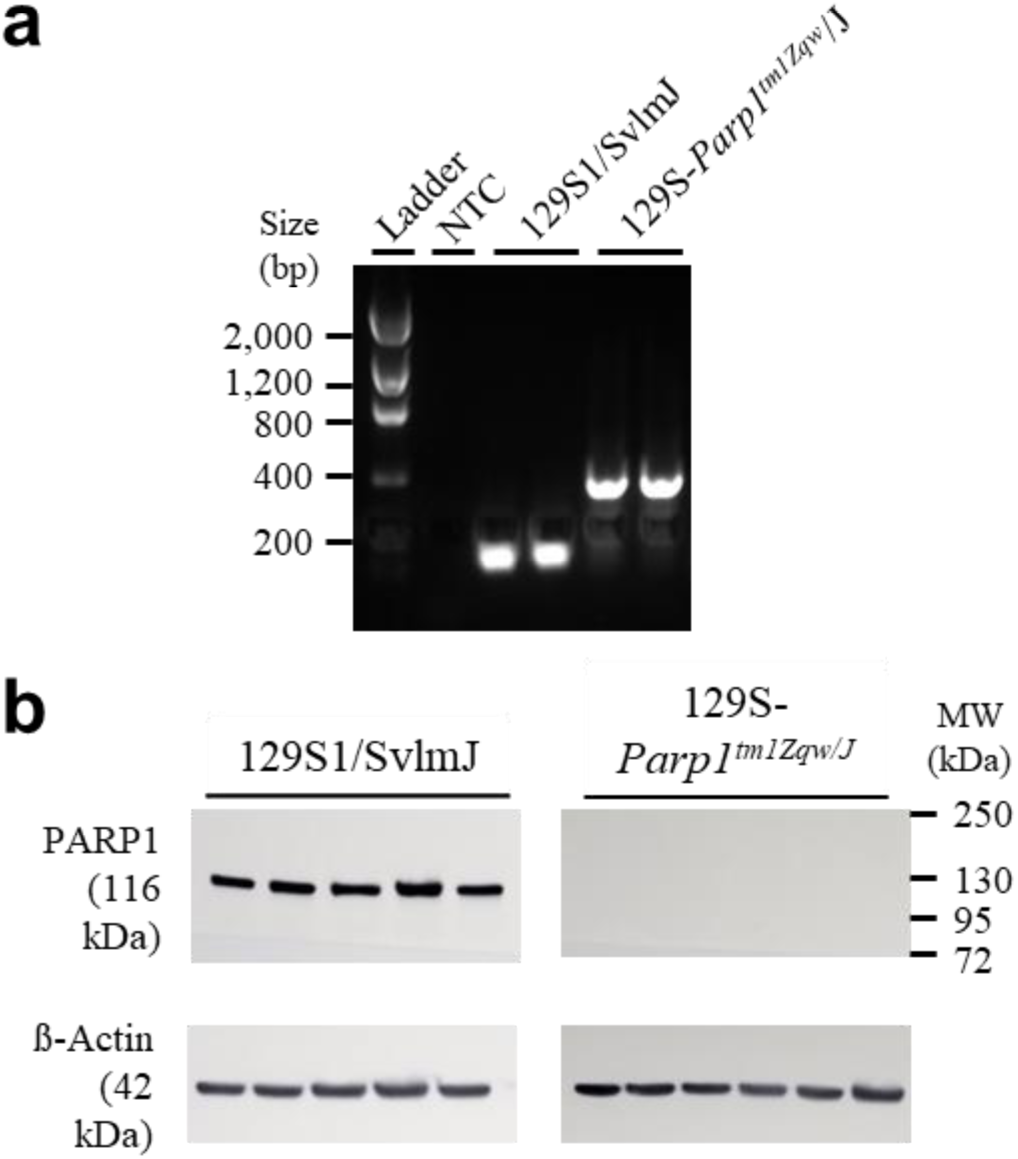
Confirmation of PARP1 deletion in 129S-*Parp1^tm1Zqw^*/J mice. (a) PARP1 genotyping PCR products of two representative 129S1/SvlmJ (WT) and two 129S-*Parp1^tm1Zqw^*/J (PARP1^-/-^) mice separated on a 1.2% agarose gel. Expected size is 112 bp for the wild type and 350 bp for the PARP1 mutant band. L, Low mass DNA ladder. NTC, no target control. (b) PARP1 Western blot of five 129S1/SvlmJ (left) and five 129S-*Parp1^tm1Zqw^*/J (right) mice demonstrating no detectable PARP1 protein (116 kDa) in the knockout mice.

**Figure S2.**
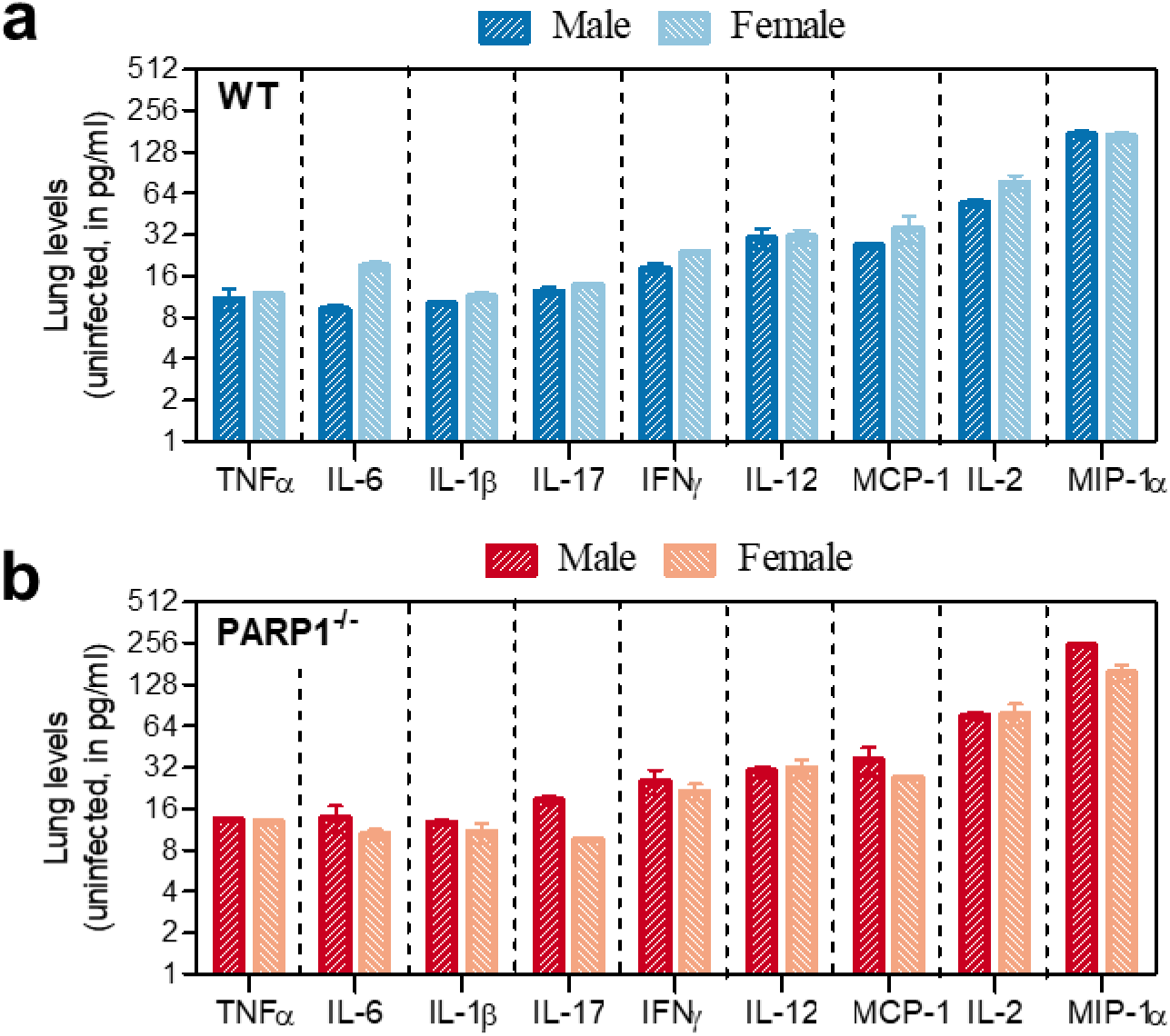
Lung levels of cytokines and chemokines in uninfected WT and PARP1^-/-^ mice do not differ by sex. Levels of lung cyto-/chemokine in uninfected male and female (a) WT or (b) PARP1^-/-^ (red) mice, quantified by Luminex multiplex assay (n=2). At baseline, cytokine and chemokine levels are unaffected by sex of PARP1 deletion.

**Figure S3.**
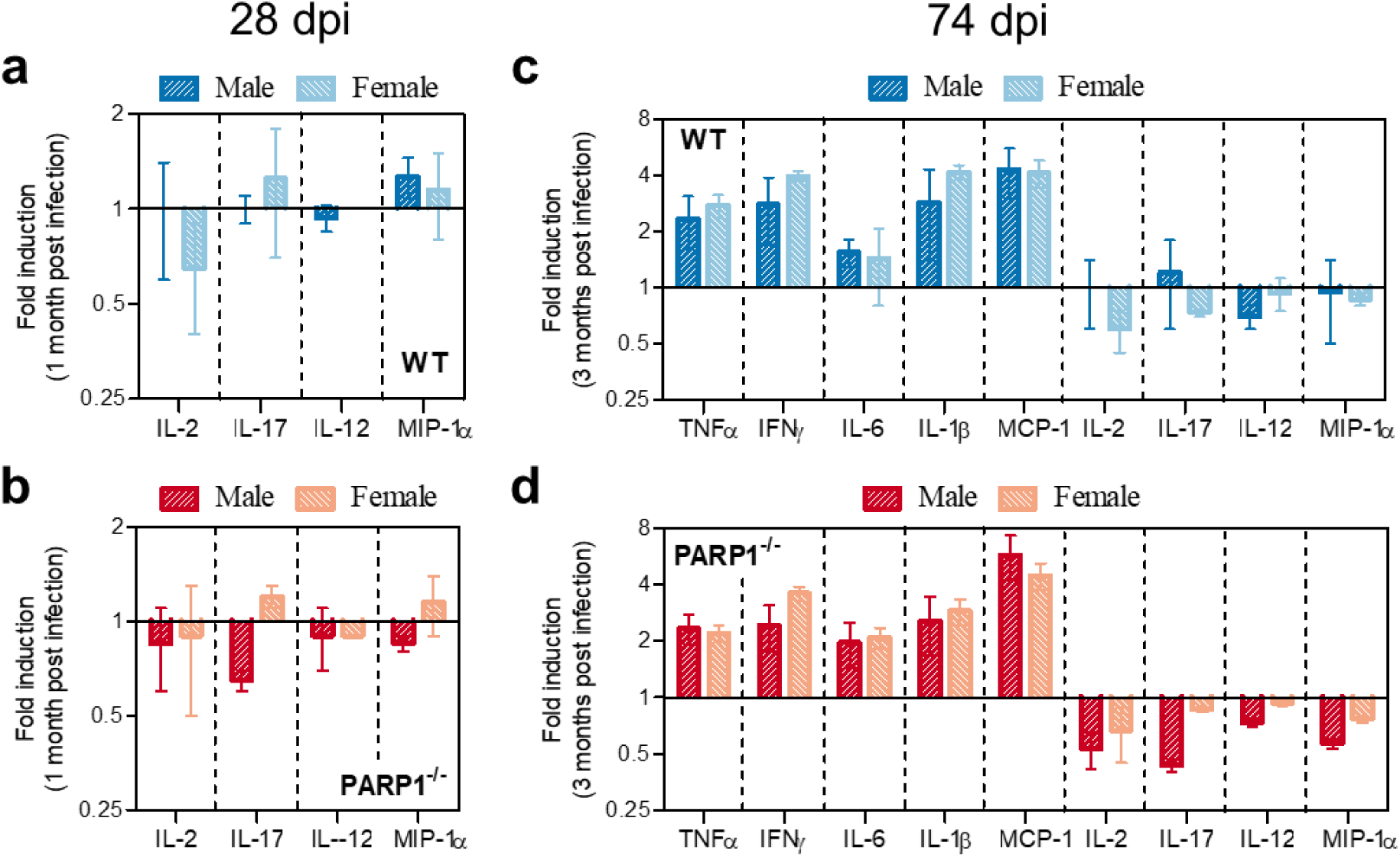
Not all TB immune responses are affected by PARP1^-/-^ status. Change in lung cyto-/chemokine levels in WT (blue) and PARP1^-/-^ (red) mice (a-b) 28 or (c-d) 74 days after aerosol infection with *M. tb* H37Rv (n=2-3). Cyto-/Chemokine levels were quantified by Luminex multiplex assay and are expressed as fold change relative to levels in uninfected lungs (set to 1; time 0, dashed line). Values above 1 indicate levels that are higher in infected than uninfected lungs whereas values less than 1 indicate levels that are reduced in infected lungs. The levels of IL-2, IL-12, IL-17 and MIP-1α were largely unaffected by *M. tb* infection, PARP1 status or sex, and at 74 dpi, most TB immune responses were unaffected by PARP1 deletion or sex.

**Figure S4.**
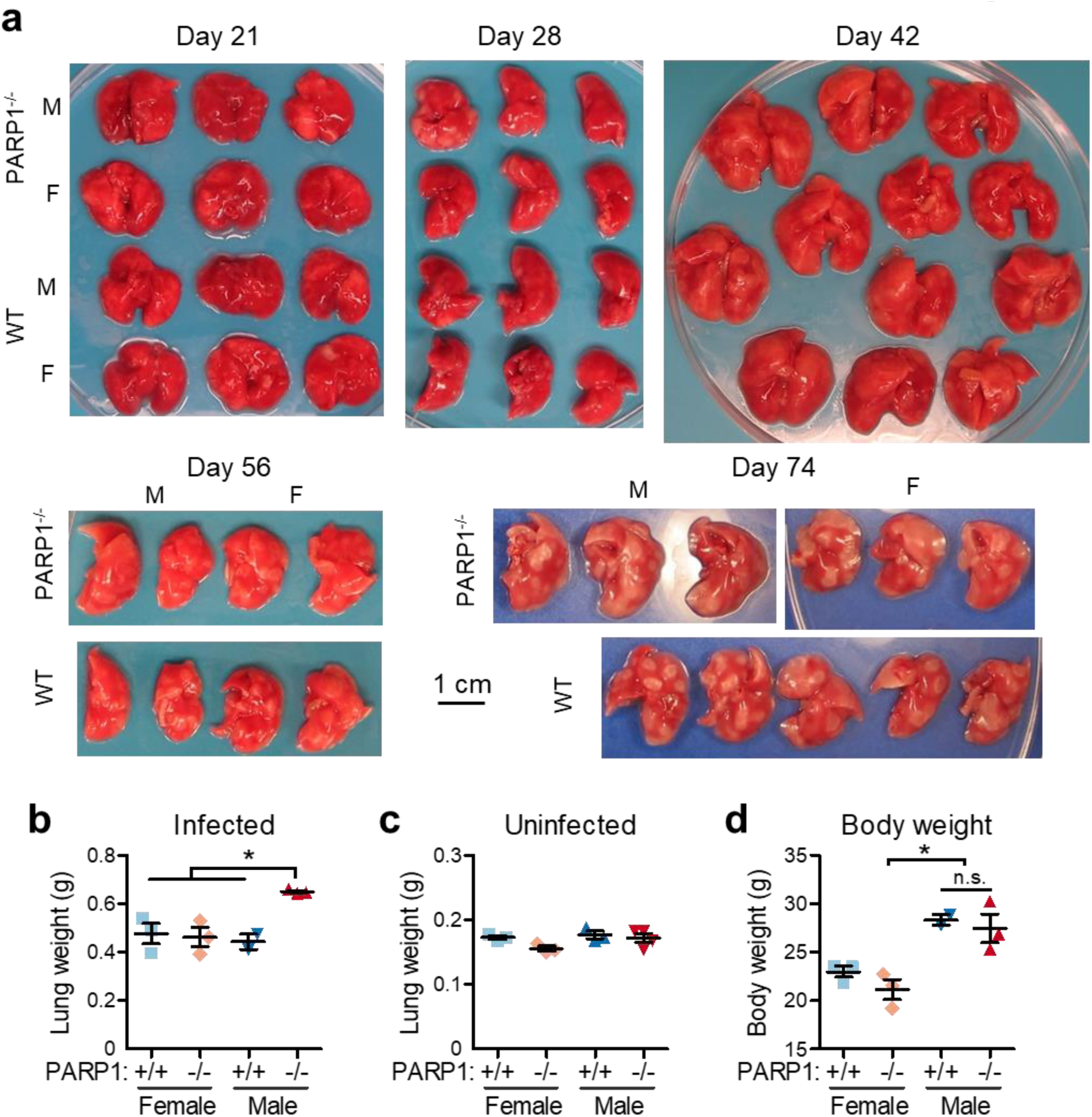
Chronically TB-infected male but not female PARP1^-/-^ mice have enlarged lungs. (a) Lung pictures of male (M) and female (F) wildtype 129S1/SvlmJ (WT) or 129S-*Parp1^tm1Zqw^*/J (PARP1^-/-^) mice at 21, 28, 42, 56 and 74 days post aerosol infection with *M. tb* H37Rv. At days 28, 56 and 74, only the left lobes (pictured) were collected for gross pathology and CFU determination and the right lobes immediately processed for cytokine and PAR level analysis. (b-d) Average weight of lungs (b-c) or mice (d) 74 days post infection or of age-matched uninfected mice (c). *, *p* < 0.05; n.s., not significant by one-way ANOVA with Bonferroni post test.

**Figure S5.**
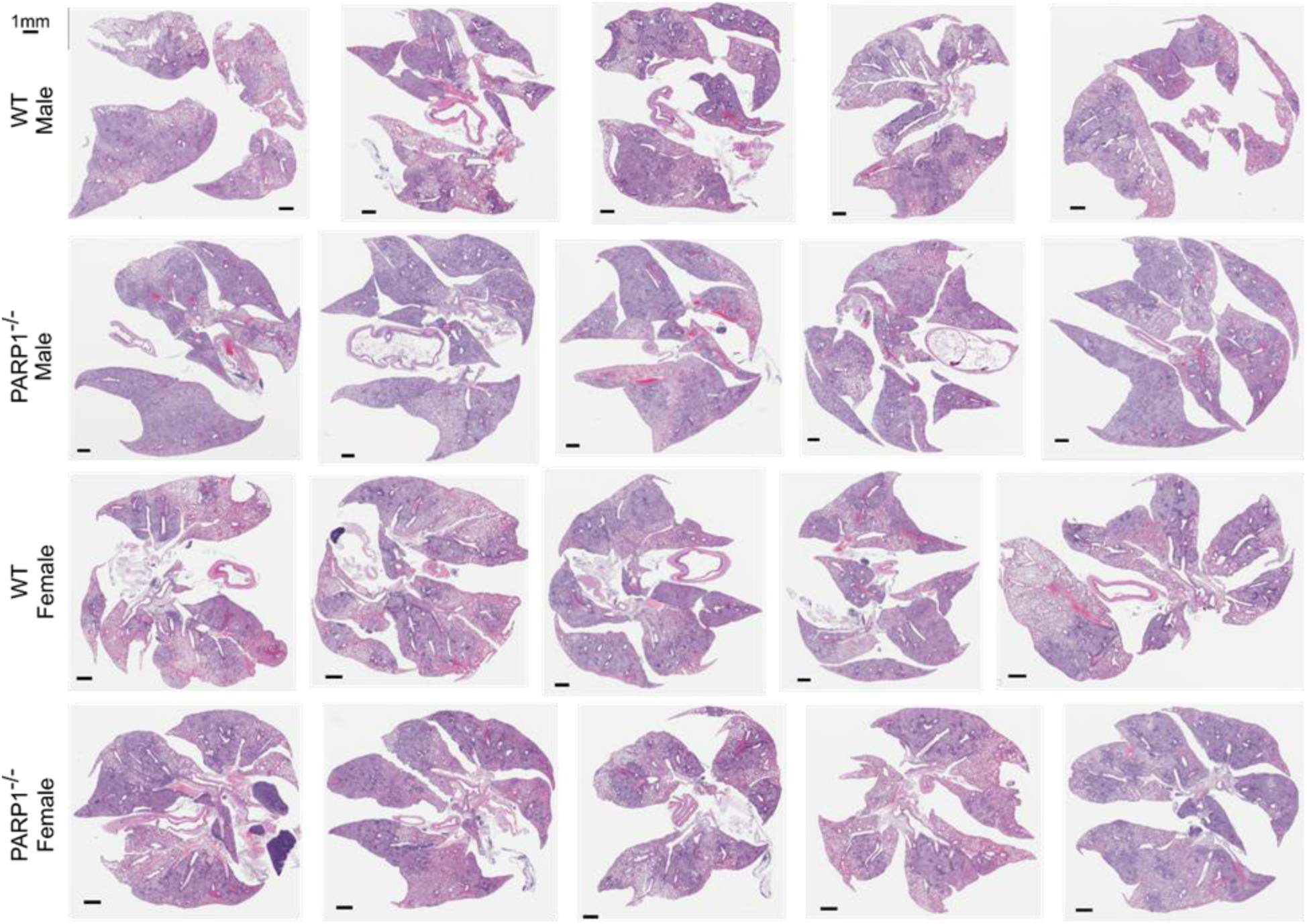
Histopathology reveals extensive lung consolidation in PARP1^-/-^ males. H&E-stained lung sections of all male and female WT or PARP1^-/-^ mice 3 months after infection with *M. tb* H37Rv (day 1: 1.66 log_10_ CFU). There was extensive consolidation throughout the lungs, in particular in PARP1^-/-^ males. Quantification of consolidation and percent lung involvement indicating lung inflammation shown in figure 4.

**Figure S6.**
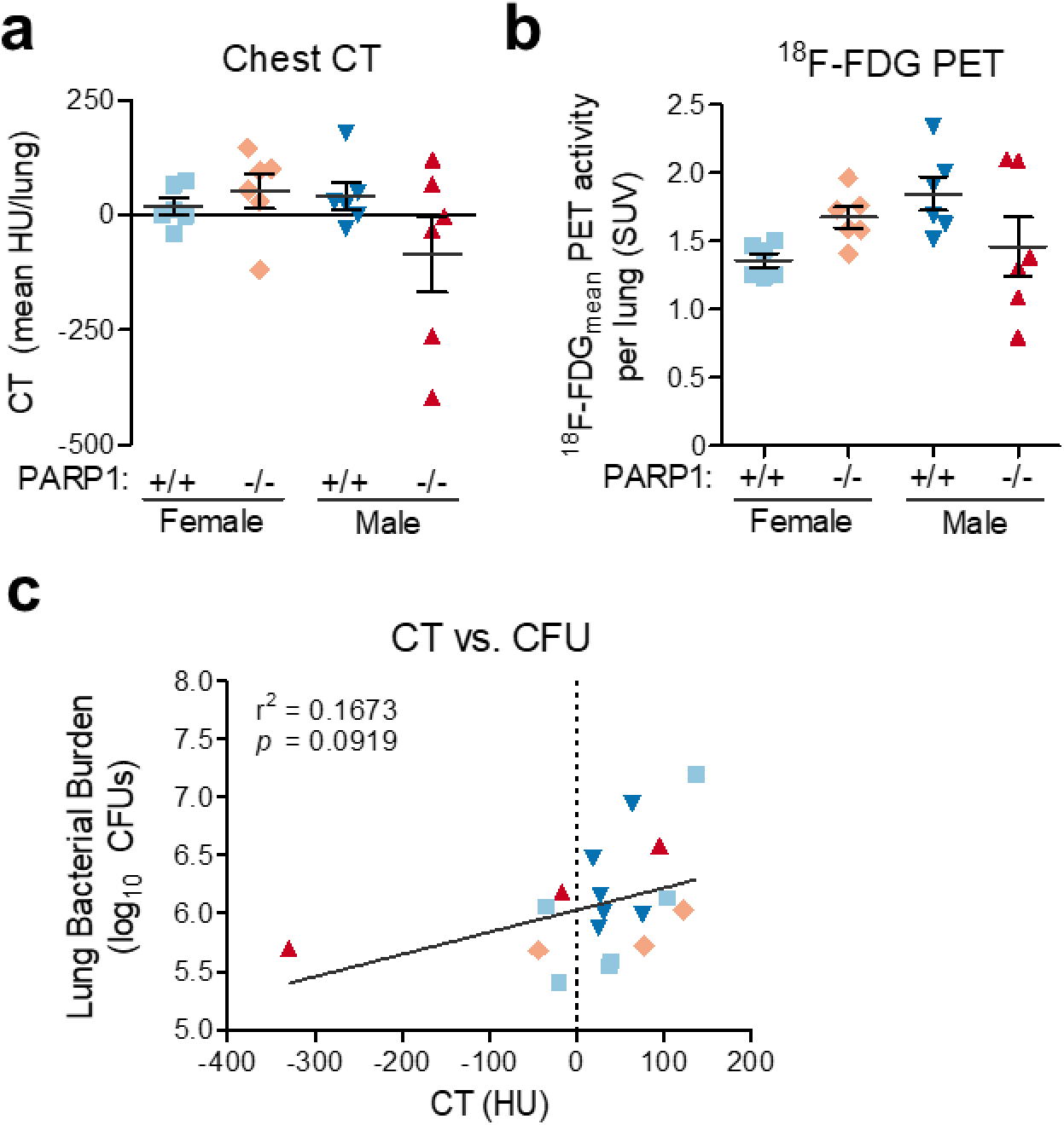
Inflammation analysis by ^18^F-FDG PET-CT. Inflammation analysis in 3 male and 3 female WT or PARP1^-/-^ mice 3 months after infection with *M. tb* H37Rv (day 1: 1.66 log_10_ CFU) by CT (a) and ^18^F-FDG PET (b). Each data point represents the mean activity recorded in the left or the right lung (2 data points per mouse). (a) CT values of −1000, 0 and +1000 HU indicate air, water and bone, respectively. CT values above 0 HU suggest diseased lung. (b) Mean ^18^F-FDG PET activity per lung was normalized to the injected dose and body weight. SUV = Standardized uptake value. (c) For each mouse, lung bacterial burden was plotted against CT (average of left and right lung). CT values correlate with bacterial burden.

**Figure S7.**
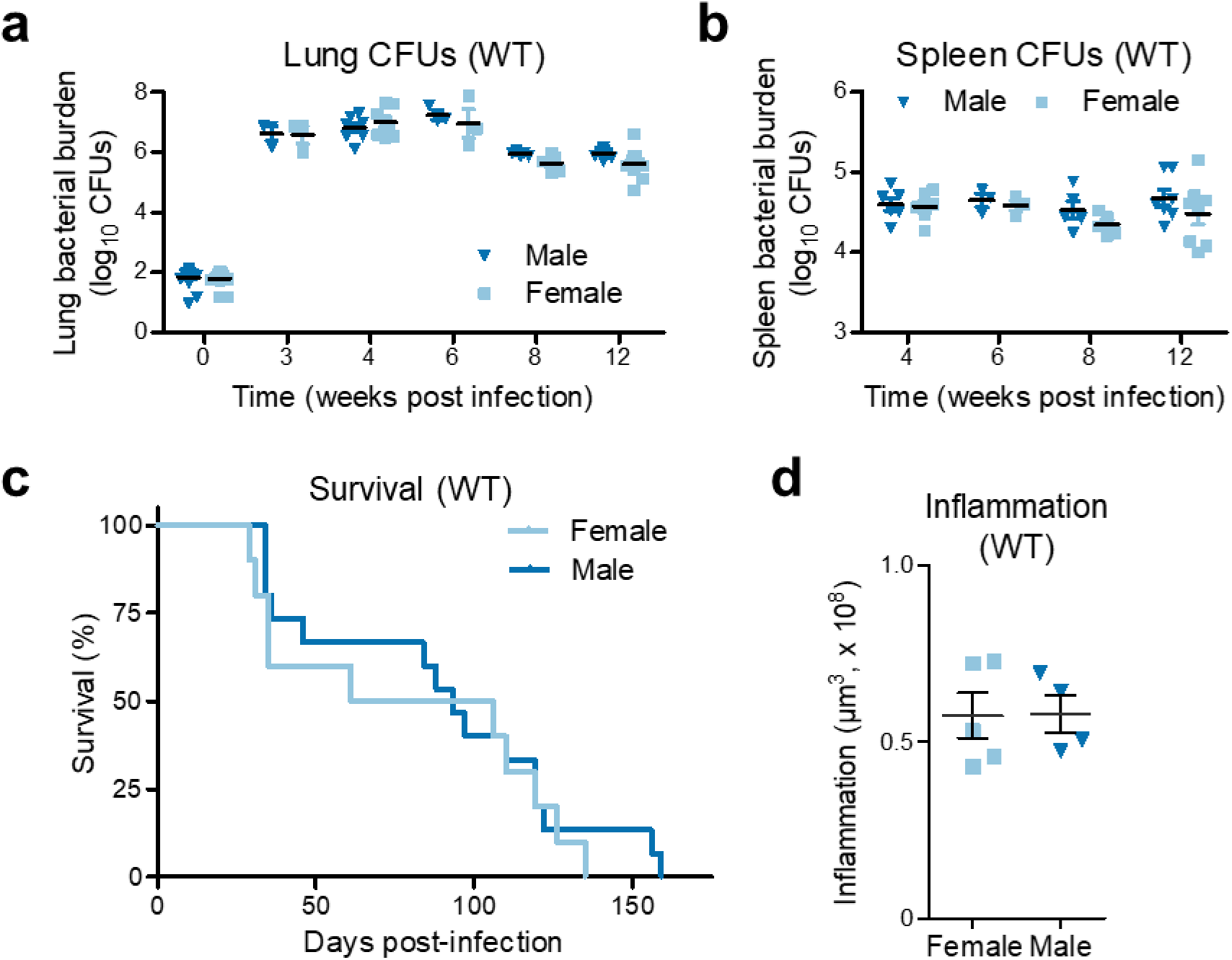
TB progression is comparable in male and female 129S1 WT mice. (**a-b**) Time course of bacterial burden in the (**a**) lungs and (**b**) spleens of male and female 129S1/SvlmJ WT mice aerosol-infected with *M. tb* H37Rv or H37RvΔ*pncA.* Data was collected in three independent experiments (day 1: 1.659 ± 0.3172, 1.724 ± 0.0916 and 1.858 ± 0.2356 log_10_ CFU (± SD)). Each data point represents an individual mouse. Lines indicate mean and SEM. (**c**) Survival in male and female 129S1/SvlmJ WT mice (n=15) following aerosol infection with *M. tb* H37Rv (day 1: 2.397 log_10_ CFU). (**d**) Lung inflammation in male and female 129S1/Svlm WT mice 3 months after aerosol infection with *M. tb* H37Rv (day 1: 1.66 log_10_ CFU). Each point represents the area of consolidation, determined histologically, in an individual mouse. There were no apparent sex differences in bacterial burden, survival or lung inflammation in TB-infected WT mice.

**Figure S8.**
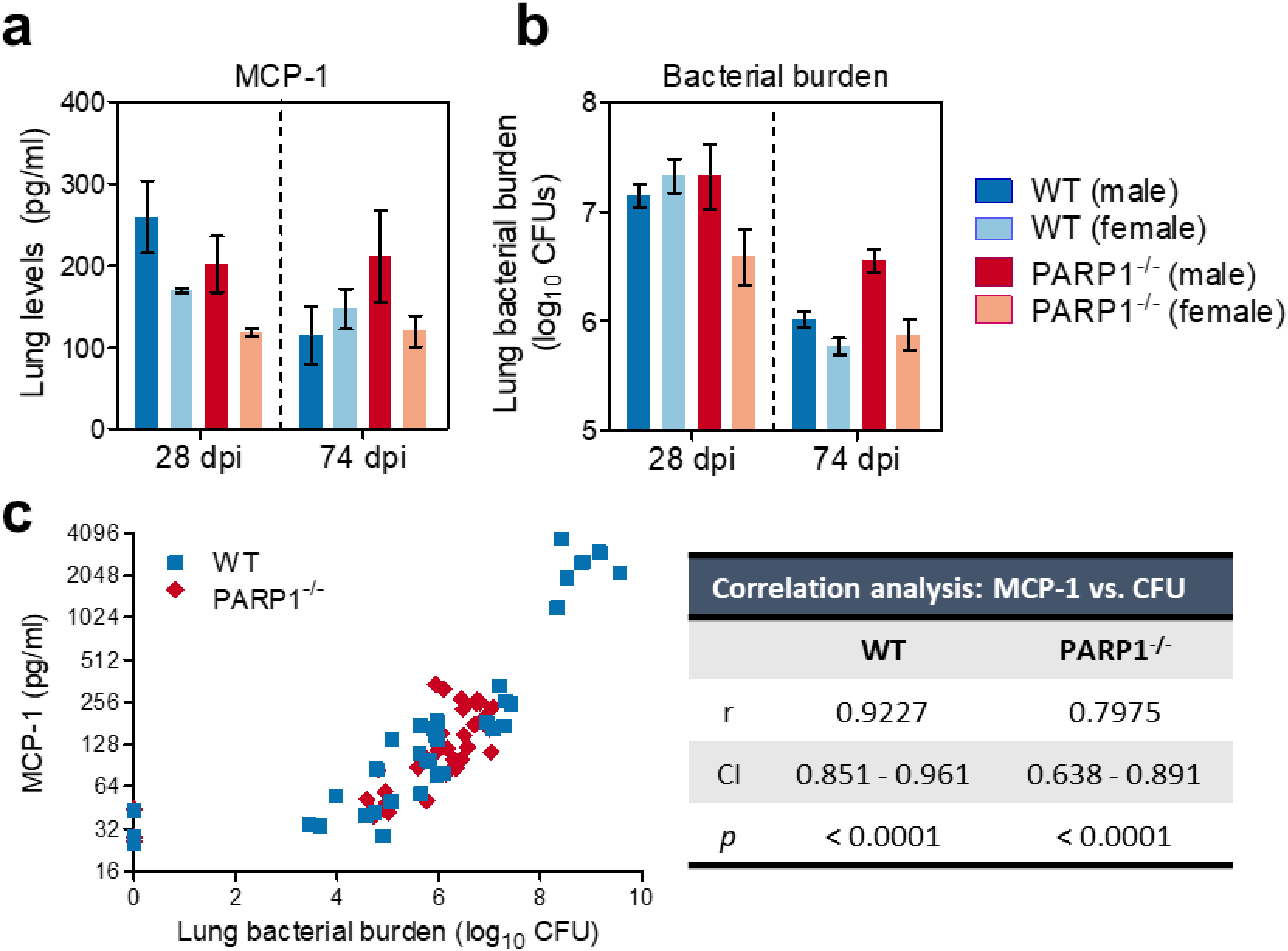
MCP-1 correlates with bacterial burden in WT and PARP1^-/-^ mice. (a-b) MCP-1 levels (a) and bacterial burden (b) in the lungs of male and female WT (blue) and PARP1^-/-^ (maroon) mice 28 ad 74 days after aerosol infection with *M. tb* H37Rv (implantation: 1.86 log_10_ CFU). Lung cytokines were quantified by Luminex multiplex assay. At 28 dpi, PARP1^-^/-females have the lowest MCP-1 levels and lowest bacterial burden, and at 74 dpi, PARP1^-^/-males have the highest MCP-1 levels and highest bacterial burden. (c) Spearman correlation analysis of bacterial burden vs. MCP-1 levels in WT (blue) and PARP1^-/-^ (red) mice. Each point represents an individual mouse. r, correlation coefficient. CI, 95% confidence interval. MCP-1 levels correlate with bacterial burden in WT and PARP1^-/-^ mice.

